# Skeletal muscle PGC-1α remodels mitochondrial phospholipidome but does not alter energy efficiency for ATP synthesis

**DOI:** 10.1101/2024.05.22.595374

**Authors:** Takuya Karasawa, Ran Hee Choi, Cesar A. Meza, J. Alan Maschek, James E. Cox, Katsuhiko Funai

## Abstract

**Background:** Exercise training is thought to improve the mitochondrial energy efficiency of skeletal muscle. Some studies suggest exercise training increases the efficiency for ATP synthesis by oxidative phosphorylation (OXPHOS), but the molecular mechanisms are unclear. We have previously shown that exercise remodels the lipid composition of mitochondrial membranes, and some of these changes could contribute to improved OXPHOS efficiency (ATP produced by O_2_ consumed or P/O). Peroxisome proliferator-activated receptor gamma coactivator-1 alpha (PGC-1α) is a transcriptional co-activator that coordinately regulates exercise-induced adaptations including mitochondria. We hypothesized that increased PGC-1α activity is sufficient to remodel mitochondrial membrane lipids and promote energy efficiency.

**Methods:** Mice with skeletal muscle-specific overexpression of PGC-1α (MCK-PGC-1α) and their wildtype littermates were used for this study. Lipid mass spectrometry and quantitative PCR were used to assess muscle mitochondrial lipid composition and their biosynthesis pathway. The abundance of OXPHOS enzymes was determined by western blot assay. High-resolution respirometry and fluorometry analysis were used to characterize mitochondrial bioenergetics (ATP production, O_2_ consumption, and P/O) for permeabilized fibers and isolated mitochondria.

**Results:** Lipidomic analyses of skeletal muscle mitochondria from wildtype and MCK-PGC-1α mice revealed that PGC-1α increases the concentrations of cone-shaped lipids such as phosphatidylethanolamine (PE), cardiolipin (CL), and lysophospholipids, while decreases the concentrations of phosphatidylcholine (PC), phosphatidylinositol (PI) and phosphatidic acid (PA). However, while PGC-1α overexpression increased the abundance of OXPHOS enzymes in skeletal muscle and the rate of O_2_ consumption (*J*O_2_), P/O values were unaffected with PGC-1α in permeabilized fibers or isolated mitochondria.

**Conclusions:** Collectively, overexpression of PGC-1α promotes the biosynthesis of mitochondrial PE and CL but neither PGC-1α nor the mitochondrial membrane lipid remodeling induced in MCK-PGC-1α mice is sufficient to increase the efficiency for mitochondrial ATP synthesis. These findings suggest that exercise training may increase OXPHOS efficiency by a PGC-1α-independent mechanism, and question the hypothesis that mitochondrial lipids directly affect OXPHOS enzymes to improve efficiency for ATP synthesis.

## Introduction

Exercise training induces a range of physiological adaptations in skeletal muscle, notably augmenting mitochondrial volume and enhancing oxidative capacity [1, 2]. There is also evidence that endurance exercise training improves the efficiency of skeletal muscle oxidative phosphorylation (OXPHOS), defined as ATP produced per oxygen consumed (P/O) [3, 4]. While mechanisms that control mitochondrial volume and respiratory function are almost exhaustively studied [5, 6], there is very little literature on the effect of exercise training on OXPHOS energy efficiency.

Peroxisome proliferator-activated receptor-γ coactivator-1α (PGC-1α) is a transcription coactivator that plays a key role in the regulation of exercise-induced mitochondrial adaptations in skeletal muscle [7–9]. Overexpression of PGC-1α in skeletal muscle robustly increases mitochondrial volume, resulting in enhanced capacity for respiration and ATP synthesis [5, 10–14]. Conversely, depletion of PGC-1α results in reduced mitochondrial volume and enzymes of OXPHOS, leading to decreased respiration and exercise capacity [15–18]. Loss-of-function experiments further indicate that PGC-1α is essential for mitochondrial cristae architecture, likely affecting bioenergetics [19]. Although no studies have examined the effect of PGC-1α on the energy efficiency of skeletal muscle OXPHOS, it is reasonable to speculate that PGC-1α may play a role in exercise training-induced improvement in skeletal muscle energy efficiency.

Mitochondria are composed of bilayer membranes that primarily consist of phospholipids. In particular, the lipid composition of the inner mitochondrial membrane (IMM) directly influences OXPHOS by modulating protein function via lipid-protein interactions, membrane properties, and cristae morphology [20]. Cone-shaped phospholipids such as phosphatidylethanolamine (PE) and cardiolipin (CL) are concentrated in the IMM presumably to induce curvatures that are found in the cristae where OXPHOS enzymes reside [21, 22]. Evidence suggests that mitochondrial PE and CL are elevated with exercise, and influence OXPHOS activity and likely efficiency [23]. Overexpression of PGC-1α increases several species of PE in skeletal muscle [24] while depletion of PGC-1α decreases total CL abundance in the heart [25]. Nonetheless, these changes represent alteration in the total cellular lipidome rather than changes that are specific to mitochondria. The purpose of the current study was to test the hypothesis that PGC-1α remodels skeletal muscle mitochondrial membrane lipid composition similarly to changes that occur with exercise training, and that these changes will coincide with improved efficiency for mitochondrial ATP synthesis (Figure 1A).

**Fig. 1.**
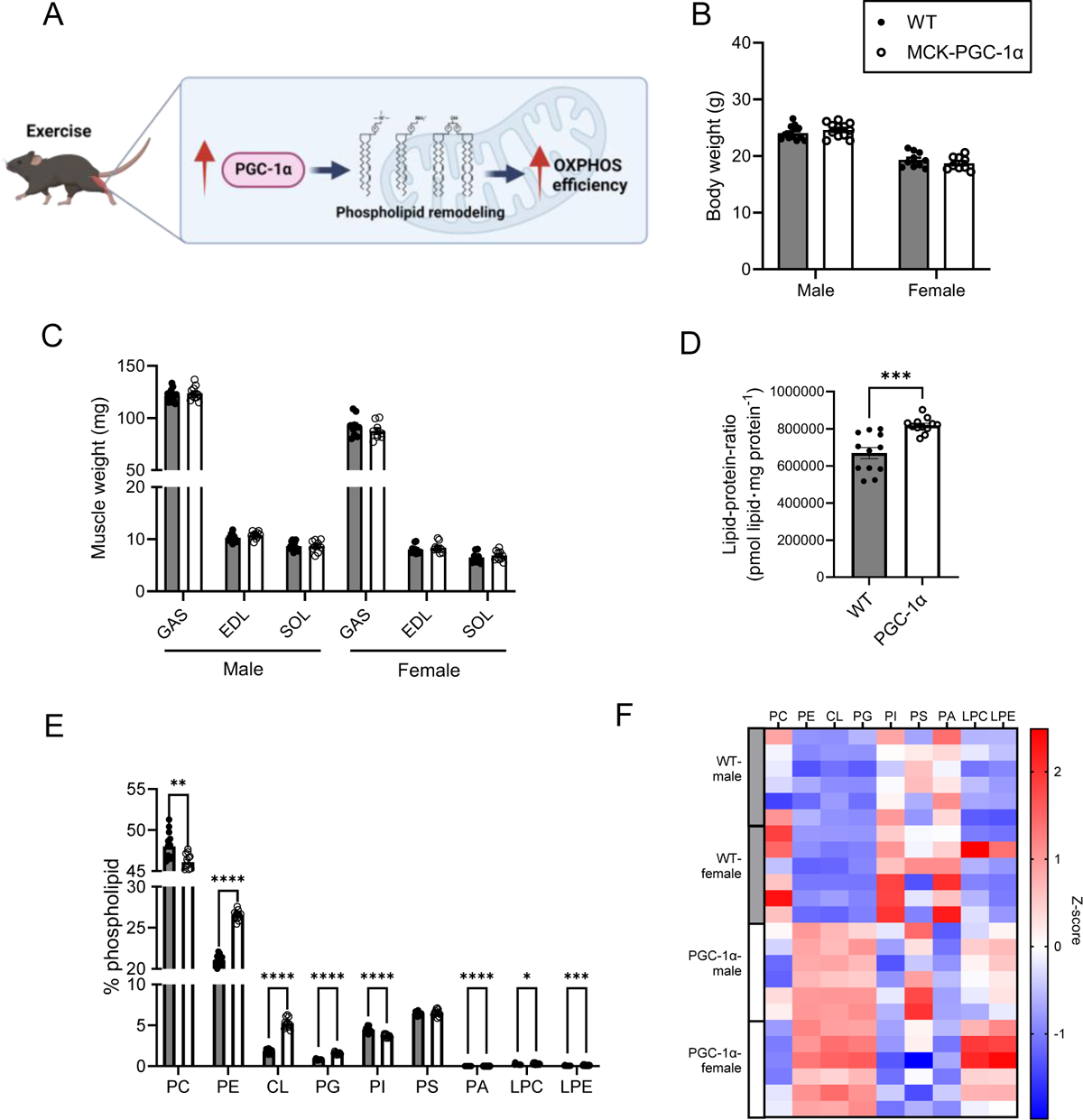
Overexpression of PGC-1α alters muscle mitochondrial phospholipid composition. (A) Schematic illustration of our hypothesis that PGC-1α increases skeletal muscle OXPHOS efficiency. (B) Body weight of male and female WT and MCK-PGC-1α mice (n = 9–14 per group). (C) Gastrocnemius (GAS), extensor digitorum longus (EDL), soleus (SOL) muscles weight (n = 9–14 per group). (D) Lipid-protein-ratio (WT n = 12, MCK-PGC-1α n = 12; combined male and female). (E) Relative abundance of each mitochondrial lipid class. (F) Heatmap of the relative abundance of each mitochondrial lipid class. Data are represented as mean ± SEM. * p<0.05, ** p<0.01, *** p<0.001, **** p<0.0001 significant difference between groups.

## Materials and methods

### Animals

Ten to twelve-week-old wildtype and MCK-PGC-1α (RRID: IMSR_JAX:008231; PGC-1α overexpression under the control of the muscle creatine kinase promoter, kindly provided by Dr. Bruce Spiegelman) littermates of both sexes were used for this study [10]. Mice were housed with a 12 h light/dark cycle in a temperature-controlled room. For terminal experiments, mice were given an intraperitoneal injection of 80 mg/kg ketamine and 10 mg/kg xylazine, after which extensor digitorum longus (EDL), soleus (SOL), and gastrocnemius (GAS) muscles were harvested. All experimental procedures were approved by the University of Utah Institutional Animal Care and Use Committee.

### Isolation of skeletal muscle mitochondria

Muscle mitochondrial isolation was performed as previously described [23]. Briefly, GAS muscle from freshly sacrificed mice was finely minced in an ice-cold mitochondrial isolation medium [MIM: 0.3 M sucrose, 10 mM 4-(2-hydroxyethyl)-1-piperazineethanesulfonic acid (HEPES), 1 mM ethylene glycol tetraacetic acid (EGTA), pH 7.1] with bovine serum albumin (BSA: 1 mg/mL). The mixture was homogenized using a TH-01 homogenizer (Omni International). Homogenates were centrifuged at 800 g for 10 min at 4°C twice. Each time, the supernatant was collected and transferred to a new tube. The supernatant was centrifuged at 12,000 g for 10 min at 4°C. The supernatant was discarded, and the pellet was resuspended in 1 mL MIM. The resuspension was centrifuged once more at 12,000 g for 10 min at 4°C, and the pellet was resuspended in 100 μL MIM.

### Mitochondrial lipid mass spectrometry

Mitochondrial phospholipids were extracted from isolated mitochondria using a modified Matyash lipid extraction protocol [26]. A mixture of ice-cold methyl-tert-butyl ether (MTBE), methanol, and internal standards [EQUISPLASH Mix (Avanti Polar Lipids 330731) and Cardiolipin Mix I (Avanti Polar Lipids LM6003)] was added to 50 μg of protein from isolated mitochondria from GAS muscle. Samples were vortexed and sonicated for 1 min before being incubated on ice for 15 min. During this time, samples were vortexed every 5 min. H_2_O was added, and the samples were again incubated on ice for 15 min with vortexing every 5 min. The samples were centrifuged at 12,000 g for 10 min. The organic (upper) layer was collected, and the aqueous layer was re-extracted with 1 mL of 10:3:2.5 (v/v/v) MTBE/ methanol/H_2_O. The MTBE layers were combined for untargeted lipidomic analysis and dried under vacuum. Lipid extracts were reconstituted in 300 μL of 8:2:2 (v/v/v) IPA/ACN/H_2_O. Lipidomics analysis was conducted at the Metabolomics Core at the University of Utah using liquid chromatography–mass spectrometry on an Agilent 6545 ultraperformance liquid chromatography-quadrupole time-of-flight mass spectrometer.

### Preparation for permeabilized muscle fiber bundles (PmFB)

PmFB were prepared as previously described [27]. Briefly, a small portion of freshly dissected red GAS muscle was placed in buffer X [7.23 mM K_2_EGTA, 2.77 mM Ca K_2_EGTA, 20 mM imidazole, 20 mM taurine, 5.7 mM ATP, 14.3 mM phosphocreatine, 6.56 mM MgCl_2_·6H_2_O, and 50 mM 2-(N-Morpholino) ethanesulfonic acid potassium salt (K-MES) (pH7.4)]. Fiber bundles were separated and permeabilized for 30 min at 4°C with 30 μg/mL saponin. After permeabilization, fiber bundles were incubated in buffer Z [105 mM K-MES, 30 mM KCl, 10 mM K_2_HPO_4_, 5 mM MgCl_2_·6H_2_O, 0.5 mg/mL BSA, and 1 mM EGTA (pH 7.4)] with 0.5 mM pyruvate and 0.2 mM malate for 15 min, and briefly washed buffer Z. PmFB samples were placed in buffer Z until analysis.

### High-resolution respirometry and fluorometry

O_2_ consumption of both the PmFB and isolated muscle mitochondria from freshly dissected GAS muscle was measured using Oroboros Oxygraph-2k (Oroboros Instruments). ATP production was assessed using Fluorolog-QM (Horiba Scientific) by enzymatically coupling ATP production to NADPH synthesis, as previously described [28]. Both experiments were conducted in buffer Z. O_2_ consumption and ATP production were stimulated by treating PmFB or isolated muscle mitochondria with 0.5 mM malate, 5 mM pyruvate, 5 mM glutamate, 10 mM succinate, and successive additions of ADP (20, 200, and 2000 μM for PmFB; 2, 20, and 200 μM for isolated muscle mitochondria). In the PmFB experiments, 20 mM creatine monohydrate and 10 μM blebbistatin were added to buffer Z to inhibit myosin adenosine triphosphatases. The P/O ratio was determined by dividing ATP production by O_2_ consumption.

### Quantitative PCR

The frozen GAS muscle was homogenized using TissueLyser II (Qiagen) in 1 ml of TRIzol (Thermo Fisher Scientific). Following homogenization, 200 μL of chloroform was added to the tube and mixed by inverting it 10 times. After 2 min, samples were centrifuged at 12,000 g for 15 min at 4°C. The aqueous supernatant was collected and placed in a new tube containing 500 μL of 100% isopropyl alcohol. The mixture was inverted a few times and incubated for 10 min. The samples were centrifuged at 12,000 g for 10 min at 4°C. The supernatant was carefully aspirated, and the pellet was washed with 75% ethyl alcohol and centrifuged at 7,500 g for 5 min. The supernatant was aspirated, and the pellet was air-dried before being resuspended in diethylpyrocarbonate-treated water. RNA was reverse-transcribed using an iScript cDNA synthesis kit (Bio-Rad). For quantitative PCR, an equal amount of cDNA was placed in a 384-well plate with SYBR Green (Thermo Fisher Scientific) and gene-specific primers. The primer sequences used are shown in Supplemental Table S1. Gene expression was analyzed using a QuantStudio 12K Flex (Life Technologies) and was normalized to ribosomal protein L32.

### Western blotting

For whole muscle tissue analysis, the frozen GAS muscle was homogenized using TH-01 homogenizer in ice-cold RIPA buffer (Thermo Fisher Scientific) supplemented with protease inhibitor cocktail (Thermo Fisher Scientific). Following homogenization, the samples were centrifuged at 12,000 g for 10 min at 4°C. The protein concentration in the supernatant was then determined using the BCA Protein Assay Kit (Thermo Fisher Scientific). For protein analysis in isolated mitochondria, the previously prepared isolated muscle mitochondria, used for high-resolution respirometry and fluorometry analysis, were utilized. Subsequent steps were identical for both sample types. Equal amounts of protein were mixed with Laemmli sample buffer (Bio-Rad) and loaded onto a 4%–20% gradient SDS-polyacrylamide gel (Bio-Rad) for electrophoresis. The proteins were then transferred from the gel to polyvinylidene fluoride membranes (Thermo Fisher Scientific). The membranes were blocked with 3% skim milk in Tris-buffered saline containing 0.1% Tween 20 (TBST) for 1 hour at room temperature, followed by overnight incubation at 4°C with primary antibodies targeting total OXPHOS (Abcam, MS604-300) or citrate synthase (Abcam, Ab96600). After washing three times with TBST, the membranes were incubated with secondary antibodies diluted 1:5,000 in 1% skim milk for 1 hour at room temperature. Following three washes in TBST and one in TBS, membranes were incubated with Western Lightning Plus-ECL (PerkinElmer) and imaged using a ChemiDoc Imaging System (Bio-Rad) and quantified with Image Lab software (Bio-Rad). The membranes were stained with Ponceau S (Sigma-Aldrich) to verify equal protein loading across lanes.

### Ex vivo skeletal muscle force production

The force production capacities of the EDL and SOL muscles were measured as previously described [29]. Briefly, both the EDL and SOL muscles were secured at their tendons, then attached to an anchor and a force transducer within a tissue bath (Aurora Scientific, Model 801C) while being submerged in oxygenated Krebs-Henseleit buffer (KHB) at 37°C. The muscles were suspended at their optimal length, determined by pulse stimulation. Subsequently, the muscles were incubated in fresh KHB and allowed to equilibrate for 5 min. Following this equilibration period, a force-frequency protocol was initiated, stimulating the muscle at increasing frequencies (10, 20, 30, 40, 60, 80, 100, 125, 150, and 200 Hz) with a 2-min rest interval between each frequency. Muscle length and mass were measured to quantify muscle cross-sectional area (CSA). Force production data were analyzed using the Aurora Scientific DMAv5.321 software, and results were reported as specific forces that normalized absolute force to muscle CSA.

### Statistical analyses

Statistical analysis was performed using GraphPad Prism 10.1.1 software. Body weight and muscle mass were analyzed separately for males and females. For analysis of all other data, both sexes were combined because no sex differences were observed in the values or the genotype effects. An unpaired Student’s t-test was used to compare differences between genotypes. For the mitochondrial bioenergetics and muscle force data, a two-way analysis of variance (ANOVA) was conducted to explore the interaction between genotype and the other factors. If a significant interaction was observed, an appropriate post hoc multiple-comparison test was subsequently applied. All data are represented as mean ± SEM, and statistical significance was set at p ≤ 0.05.

## Results

### PGC-1α overexpression alters muscle mitochondrial lipidome

Wildtype and MCK-PGC-1α mice did not differ in body weights and muscle weights (Figure 1B,C). Lipidomics analyses revealed a number of robust changes in the skeletal muscle mitochondrial lipidome. First, PGC-1α overexpression promoted a 22% increase in lipid-to-protein ratio (Figure 1D), suggesting that PGC-1α increases mitochondrial membrane lipid syntheses to a greater extent than mitochondrial proteins. PGC-1α overexpression increased the concentrations of PE, CL, and its precursor phosphatidylglycerol (PG), as well as lysophosphatidylcholine (LPC) and lysophosphatidylethanolamine (LPE) (Figure 1E,F).

### Skeletal muscle PGC-1α overexpression promotes PE and CL biosynthesis in isolated mitochondria

By their relative abundance, PGC-1α overexpression has the most robust effects to increase PE and CL concentrations in mitochondria compared to other lipid classes. For both PE and CL, however, abundance of some species was reduced with PGC-1α overexpression (Figure 2A, Figure S1). With respect to PE, PGC-1α overexpression tended to reduce species containing 16-carbon acyl chains in the SN1 position (Figure 2A,B). In the CL class, PGC-1α led to a substantial increase in TLCL (tetralinoleoyl-CL or 18:2/18:2/18:2/18:2-CL, considered to be a functional form of CL) abundance, resulting in nearly a twofold higher ratio of TLCL to total CL compared to WT mice (Figure 2A,C). CL of low molecular weights also tended to be lower in female mice vs. male mice, regardless of genotype.

**Fig. 2.**
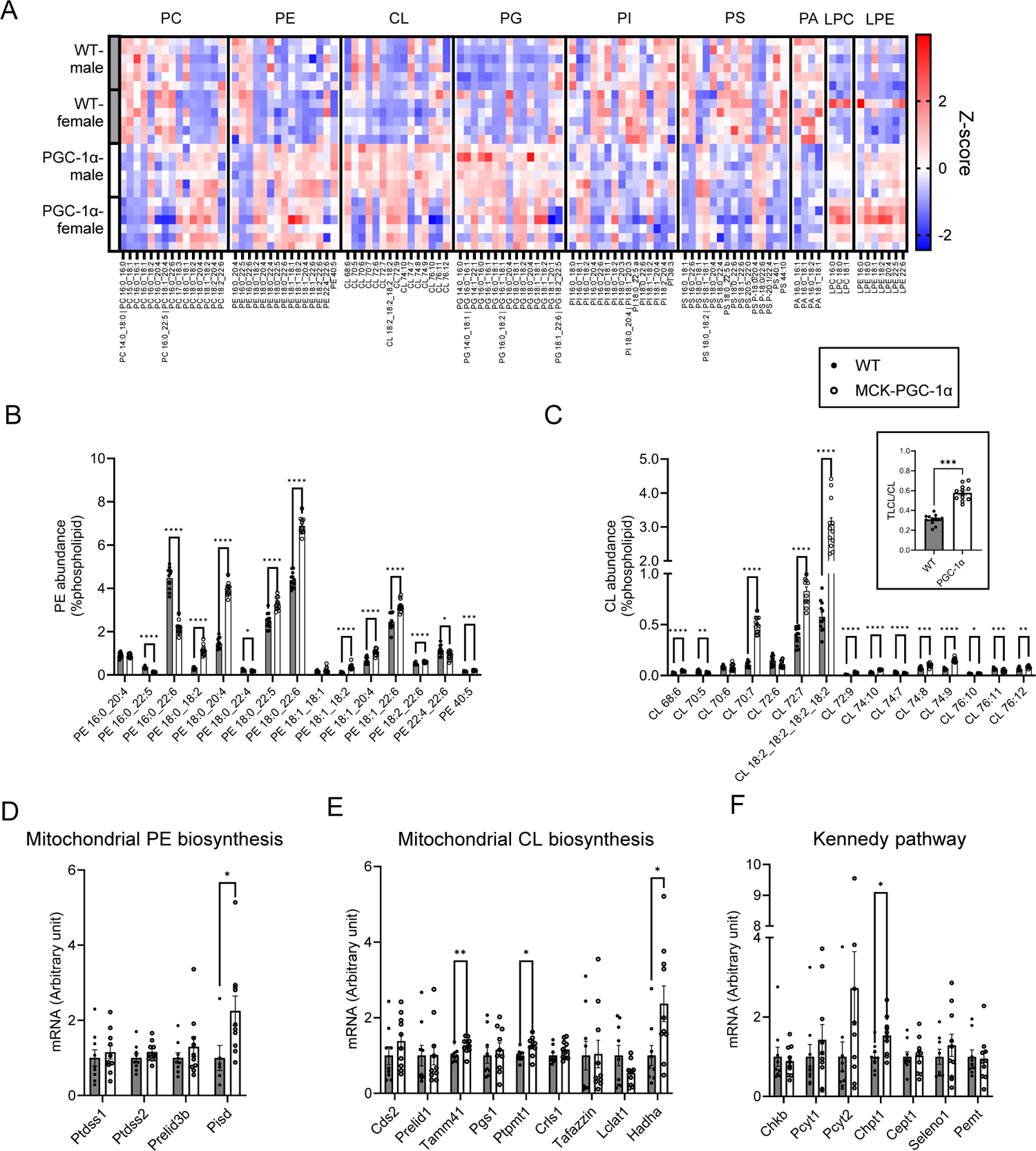
PGC-1α promotes PE and CL biosynthesis in isolated muscle mitochondria. (A) Heatmap of the relative abundance of mitochondrial lipids. The X-axis represents individual lipid species classified according to lipid classes (WT n = 12, MCK-PGC-1α n = 12). (B) Abundance of individual phosphatidylethanolamine (PE) species in isolated mitochondria. (C) Abundance of individual cardiolipin (CL) species in isolated mitochondria. The ratio of tetralinoleoyl CL (TLCL) to total CL is presented as an insert. (D–F) mRNA expression of enzymes and transporters that relate to biosynthesis of mitochondrial PE (D) and CL (E), and Kennedy pathway (F) (WT n = 9, MCK-PGC-1α n = 10). Data are combined male and female results and are represented as mean ± SEM. * p<0.05, ** p<0.01, *** p<0.001, **** p<0.0001 significant difference between groups.

Because PGC-1α altered PE and CL abundance, we examined whether PGC-1α influenced the gene expression of enzymes involved in mitochondrial PE and CL biosynthesis. Mitochondrial PE is primarily generated from phosphatidylserine (PS) by the PS decarboxylase (Pisd) localized to the IMM, which was increased by PGC-1α overexpression (Figure 2D). CL is almost exclusively synthesized by a series of enzymes localized in the IMM that convert PA into PG, PG into CL, and convert generic CL into TLCL. PGC-1α increased or tended to increase mRNA levels for some of these enzymes (Figure 2E). There were additional modest and nuanced changes in the mRNA levels for enzymes in the endoplasmic reticulum phospholipid biosynthesis (Kennedy pathway) that supplies substrates for mitochondrial lipid biosynthesis (Figure 2F).

### PGC-1α overexpression enhances respiration capacity but does not change OXPHOS efficiency in PmFB

Consistent with previous reports [10, 14], PGC-1α overexpression increased the abundance of OXPHOS subunits and citrate synthase in whole muscle lysates (Figure 3A,B). High-resolution respirometry demonstrated that PGC-1α overexpression increases O_2_ consumption (*J*O_2_) in permeabilized fibers (PmFB) (Figure 3C). PGC-1α also tended to increase the ATP production rate (*J*ATP), although to a lesser extent than *J*O_2_ (Figure 3D). Together, PGC-1α did not alter P/O in skeletal muscle PmFB (the mean was lower, not higher, in MCK-PGC-1α compared to wildtype) (Figure 3E).

**Fig. 3.**
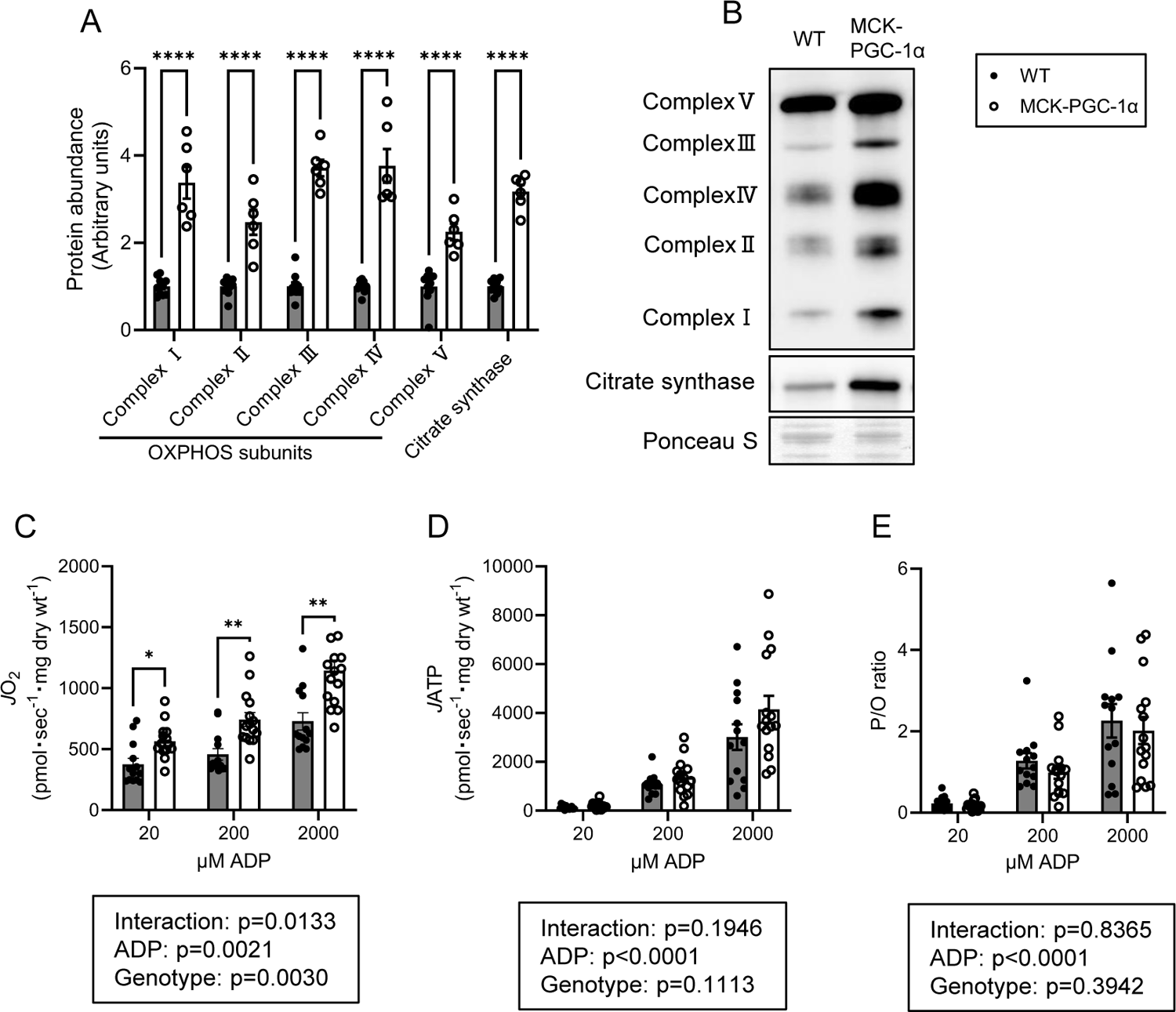
PGC-1α overexpression does not change OXPHOS energy efficiency in PmFB. (A) Abundance of OXPHOS subunits and citrate synthase in the whole lysate of gastrocnemius muscle (WT n = 10, MCK-PGC-1α n = 6). (B) Representative western blots of OXPHOS subunits and citrate synthase. (C–E) Rate of ATP production (C), rate of oxygen consumption (D), and P/O ratio (E) in fiber bundles isolated from gastrocnemius muscle (WT n = 13, MCK-PGC-1α n = 15). Data are combined male and female results and are represented as mean ± SEM. *p<0.05, ** p<0.01, **** p<0.0001 significant difference between groups.

### PGC-1α overexpression does not robustly alter the OXPHOS efficiency in isolated muscle mitochondria

While studying PmFB allows us to examine mitochondria in an ecosystem that is more similar to the in vivo environment, the preparation procedures also yield organelles and enzyme systems that may interfere with mitochondrial metabolism in situ. Thus, to further study the influence of PGC-1α on OXPHOS efficiency, we performed the same set of experiments also in isolated muscle mitochondria. Because units are normalized to the abundance of mitochondrial proteins, the effect of PGC-1α to increase OXPHOS subunits (Figure 3A,B) largely went away except for modest increases in subunits for Complexes III and IV (Figure 4A,B). Contrary to our hypothesis, PGC-1α overexpression tended to reduce *J*O_2_, *J*ATP, and P/O in skeletal muscle mitochondria (Figure 4C–E).

**Fig. 4.**
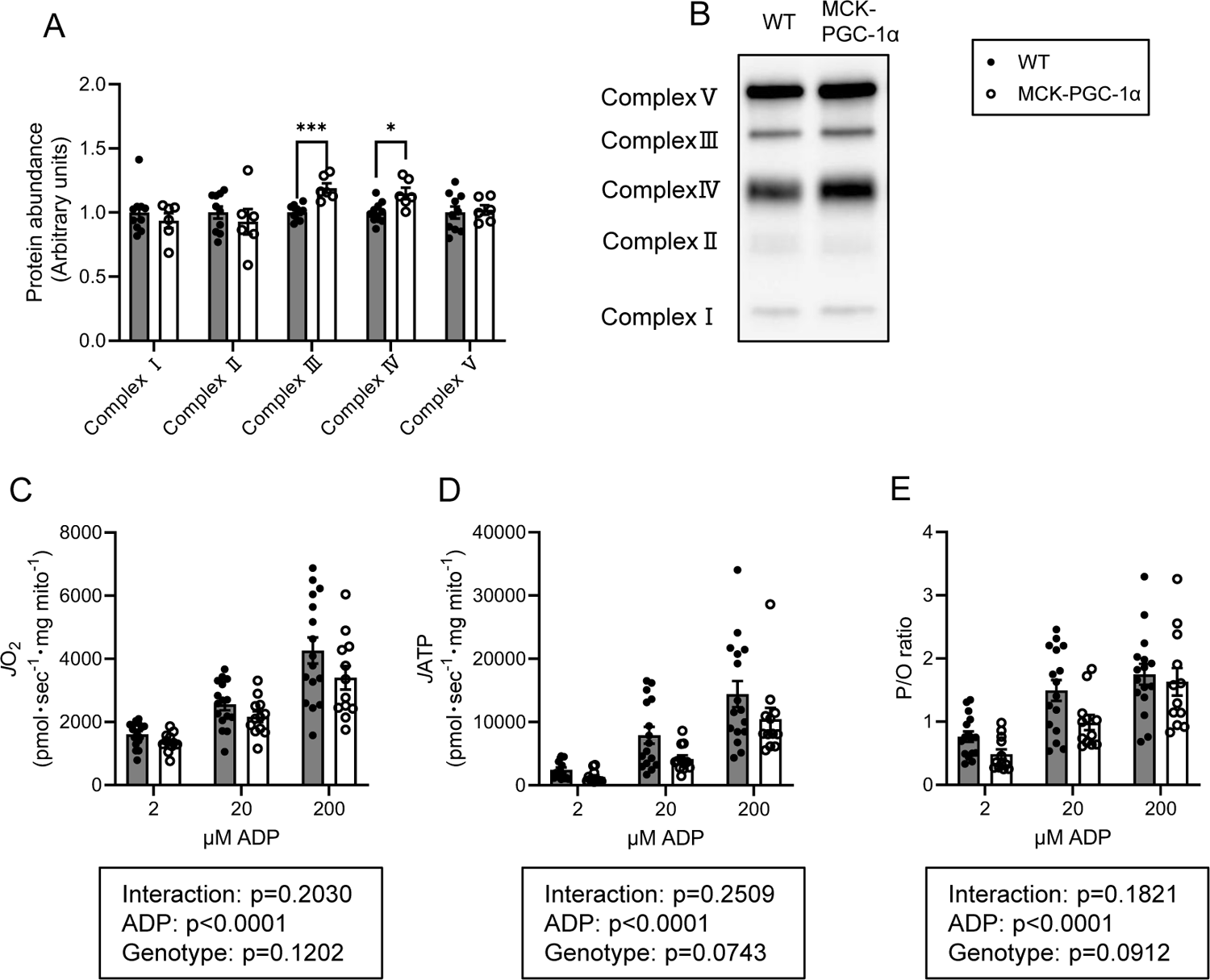
PGC-1α overexpression does not alter OXPHOS energy efficiency in isolated muscle mitochondria. (A) Abundance of OXPHOS subunits in isolated mitochondria from gastrocnemius muscle (WT n = 10, MCK-PGC-1α n = 6). (B) Representative western blots of OXPHOS subunits. (C–E) Rate of ATP production (C), rate of oxygen consumption (D), and P/O ratio (E) in isolated mitochondria from gastrocnemius muscle (WT n = 13, MCK-PGC-1α n = 15). Data are combined male and female results and are represented as mean ± SEM. * p<0.05, *** p<0.001 significant difference between groups.

### Skeletal muscle PGC-1α overexpression reduces force-generating capacity

Prompted by trends for reduced *J*ATP (p=0.0743) and P/O (p=0.0912), we examined a potential functional consequence of such bioenergetic shift by PGC-1α overexpression. Surprisingly, but consistent with the results from high-resolution fluorometry and respirometry, PGC-1α overexpression reduced the force-generating capacity of fast-twitch EDL muscle (Figure 5A,B) although the reduction was less pronounced in the slow-twitch SOL muscle (Figure 5C,D).

**Fig. 5.**
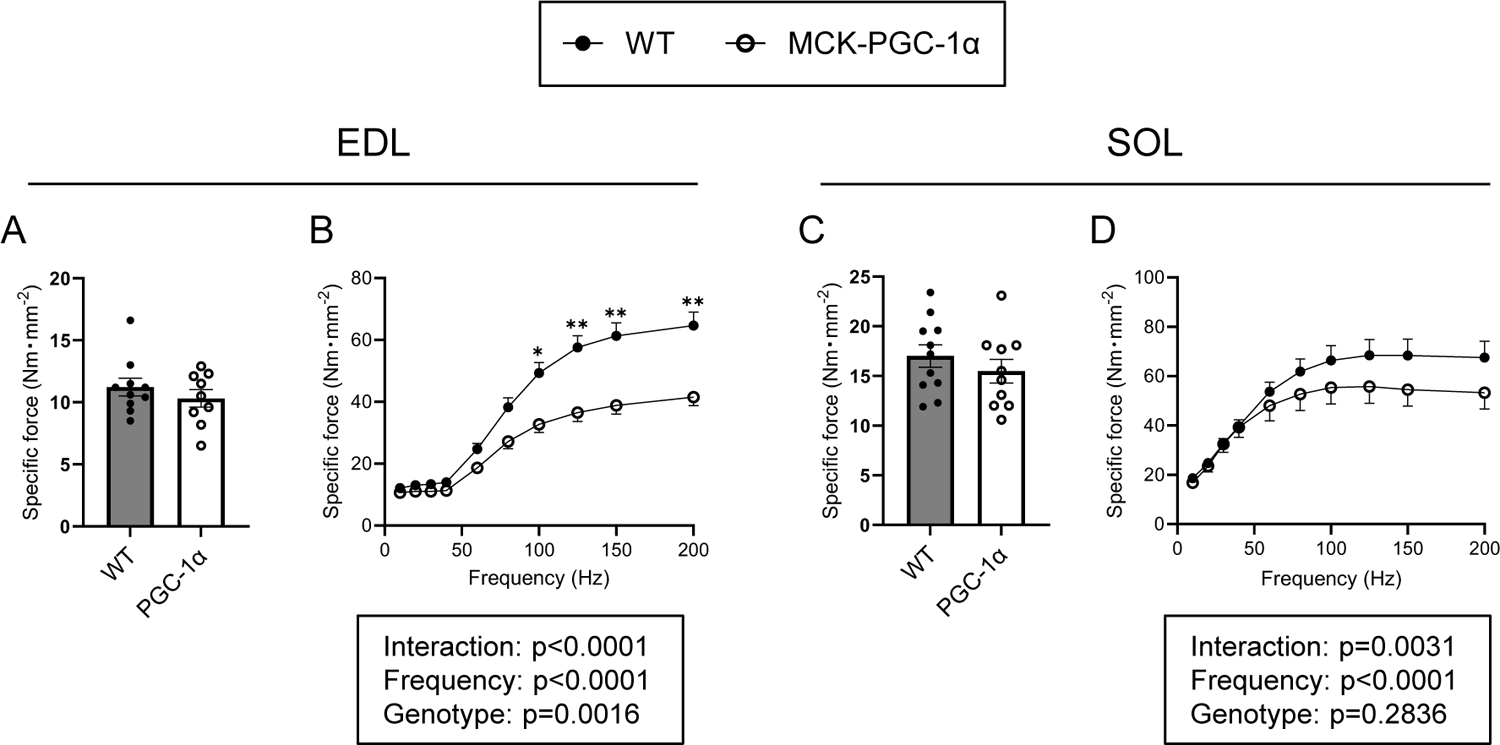
Overexpression of PGC-1α reduces muscle force production in EDL muscle. (A,B) Force produced with a pulse stimulation (A) and tetanic stimulation ranging from 10 to 200 Hz (B) in extensor digitorum longus (EDL) muscle (n = 9–11). (C,D) Force produced with a pulse stimulation (B) and tetanic stimulation ranging from 10 to 200 Hz (D) in soleus (SOL) muscle (n = 9–11 per group). Data are combined male and female results and are represented as mean ± SEM. *p<0.05, ** p<0.01 significant difference between groups.

## Discussion

Exercise training appears to increase the efficiency for ATP synthesis by oxidative phosphorylation (OXPHOS) [3, 4]. This study was conducted to test the hypothesis that: 1) PGC-1α remodels skeletal muscle mitochondrial membrane lipid composition similarly to changes that occur with exercise training, and 2) such change may mediate an increase in the energy efficiency for mitochondrial ATP synthesis. PGC-1α overexpression indeed induced remodeling of the mitochondrial membrane lipids, primarily for the biosynthesis of mitochondrial PE and CL. However, PGC-1α overexpression alone was insufficient to increase mitochondrial P/O, suggesting that additional mechanisms are essential for exercise-induced increase in OXPHOS energy efficiency.

It is well-documented that exercise training promotes mitochondrial biogenesis and increases oxidative capacity [1, 2]. However, less is known about training-induced adaptations in efficiency for mitochondrial ATP synthesis, that in turn could also influence oxidative capacity. Studies in rats and humans have demonstrated an increase in efficiency for mitochondrial OXPHOS following training. Skeletal muscle mitochondria from rats subjected to 8 weeks of running exercise training demonstrated higher OXPHOS energetic efficiency compared to sedentary rats [3]. In humans, 6 weeks of high-intensity training enhanced was found to improve skeletal muscle mitochondrial OXPHOS efficiency concomitant to greater performance efficiency [4]. To our knowledge, mechanisms that mediate exercise-induced increase in mitochondrial energy efficiency had not been previously reported.

We have previously shown that exercise remodels the lipid composition of mitochondrial membranes that in turn influences mitochondrial respiratory enzymes [23]. Given the well-known role of PGC-1α in mediating many of the exercise responses in skeletal muscle, we initially hypothesized that PGC-1α remodels skeletal muscle mitochondrial membrane lipid composition that in turn increases OXPHOS efficiency. Indeed, PGC-1α overexpression promoted robust changes in mitochondrial membrane lipid position including increased PE and CL, changes that resembled those that occur with exercise training as well as differences observed between high-capacity running rats and low-capacity running rats [23]. PGC-1α overexpression also increased the proportion of TLCL as seen in overloaded mouse skeletal muscle [14]. These findings are consistent with the notion that PGC-1α mediates the exercise-induced mitochondrial membrane lipid modeling that occurs in skeletal muscle.

We previously demonstrated that exercise training increased Pisd expression in mouse skeletal muscle [23]. Given that overexpression of PGC-1α significantly increased Pisd mRNA level, it can be deduced that PGC-1α likely mediates exercise-induced increase in skeletal muscle Pisd expression. We also observed increases in mRNA levels of enzymes involved in TLCL biosynthesis. Prola et al. demonstrated that reduced CL coincides with lower muscle P/O, and reconstitution of mitochondrial CL with unilamellar vesicles can restore P/O [30]. In our previous study, we demonstrated that weight loss also increases TLCL and CL transacylase tafazzin [27]. In that study, we also demonstrated that global tafazzin knockdown decreases muscle P/O and increases systemic metabolic rate. It is unclear why PGC-1α-induced increase in TLCL did not coincide with improved muscle P/O. A plausible scenario is that an increase in TLCL is necessary, but not sufficient for exercise-induced increase in P/O. Our data suggest that such additional mechanisms that contribute to exercise-induced increase in P/O are likely PGC-1α independent.

Likely as one of the consequences for the reduced energy efficiency, MCK-PGC-1α overexpression decreased maximal force in the EDL muscles. This study is not the first to report PGC-1α overexpression attenuated muscle force production [14, 31]. Summermatter et al. suggested that overexpression of PGC-1α decreases calcium release contributing to reduced maximal force [31]. Our data suggest a potential bioenergetic component to lower force-generating capacity.

In conclusion, the current study shows that overexpression of PGC-1α does not alter OXPHOS efficiency in skeletal muscle. While overexpression of PGC-1α promoted the biosynthesis of mitochondrial PE and CL and increased the abundance of OXPHOS enzymes in skeletal muscle, these changes were insufficient to increase P/O such that occurs with exercise training. These findings suggest that exercise training promotes additional PGC-1α-independent mechanisms that contribute to increased OXPHOS efficiency. There remains a possibility that these mechanisms synergize with PGC-1α promoted mechanisms (including mitochondrial membrane lipid remodeling) to increase skeletal muscle mitochondrial P/O.

## Acknowledgments

This research is supported by an NIH grant (DK107397, GM144613, DK197397, and AG074535 to K.F.) and the Grant-in-aid for Japan Society for Promotion of Science (JSPS) Fellows (22J15092 to T.K.). University of Utah Metabolomics Core Facility is supported by S10 OD016232, S10 OD021505, and U54 DK110858. We would like to thank Drs. Bruce Spiegelman and Melanie Mittenbühler (Harvard Medical School, Boston, MA, USA) for providing the MCK-PGC-1α mice.

## References

1. Holloszy JO. Biochemical adaptations in muscle. Effects of exercise on mitochondrial oxygen uptake and respiratory enzyme activity in skeletal muscle. J Biol Chem. 1967;242:2278–82.

2. Henriksson J, Reitman JS. Time course of changes in human skeletal muscle succinate dehydrogenase and cytochrome oxidase activities and maximal oxygen uptake with physical activity and inactivity. Acta Physiol Scand. 1977;99:91–7.

3. Zoladz JA, Koziel A, Woyda-Ploszczyca A, Celichowski J, Jarmuszkiewicz W. Endurance training increases the efficiency of rat skeletal muscle mitochondria. Pflugers Arch. 2016;468:1709–24.

4. Fiorenza M, Lemminger AK, Marker M, Eibye K, Iaia FM, Bangsbo J, et al. High-intensity exercise training enhances mitochondrial oxidative phosphorylation efficiency in a temperature-dependent manner in human skeletal muscle: implications for exercise performance. FASEB J. 2019;33:8976–89.

5. Wu Z, Puigserver P, Andersson U, Zhang C, Adelmant G, Mootha V, et al. Mechanisms controlling mitochondrial biogenesis and respiration through the thermogenic coactivator PGC-1. Cell. 1999;98:115–24.

6. Egan B, Zierath JR. Exercise metabolism and the molecular regulation of skeletal muscle adaptation. Cell Metab. 2013;17:162–84.

7. Baar K, Wende AR, Jones TE, Marison M, Nolte LA, Chen M, et al. Adaptations of skeletal muscle to exercise: rapid increase in the transcriptional coactivator PGC-1. FASEB J. 2002;16:1879–86.

8. Terada S, Goto M, Kato M, Kawanaka K, Shimokawa T, Tabata I. Effects of low-intensity prolonged exercise on PGC-1 mRNA expression in rat epitrochlearis muscle. Biochem Biophys Res Commun. 2002;296:350–4.

9. Wright DC, Han DH, Garcia-Roves PM, Geiger PC, Jones TE, Holloszy JO. Exercise-induced mitochondrial biogenesis begins before the increase in muscle PGC-1alpha expression. J Biol Chem. 2007;282:194–9.

10. Lin J, Wu H, Tarr PT, Zhang CY, Wu Z, Boss O, et al. Transcriptional co-activator PGC-1 alpha drives the formation of slow-twitch muscle fibres. Nature. 2002;418:797–801.

11. Choi CS, Befroy DE, Codella R, Kim S, Reznick RM, Hwang YJ, et al. Paradoxical effects of increased expression of PGC-1alpha on muscle mitochondrial function and insulin-stimulated muscle glucose metabolism. Proc Natl Acad Sci U S A. 2008;105:19926–31.

12. Calvo JA, Daniels TG, Wang X, Paul A, Lin J, Spiegelman BM, et al. Muscle-specific expression of PPARgamma coactivator-1alpha improves exercise performance and increases peak oxygen uptake. J Appl Physiol (1985). 2008;104:1304–12.

13. Hoeks J, Arany Z, Phielix E, Moonen-Kornips E, Hesselink MK, Schrauwen P. Enhanced lipid-but not carbohydrate-supported mitochondrial respiration in skeletal muscle of PGC-1alpha overexpressing mice. J Cell Physiol. 2012;227:1026–33.

14. Perez-Schindler J, Summermatter S, Santos G, Zorzato F, Handschin C. The transcriptional coactivator PGC-1alpha is dispensable for chronic overload-induced skeletal muscle hypertrophy and metabolic remodeling. Proc Natl Acad Sci U S A. 2013;110:20314–9.

15. Adhihetty PJ, Uguccioni G, Leick L, Hidalgo J, Pilegaard H, Hood DA. The role of PGC-1alpha on mitochondrial function and apoptotic susceptibility in muscle. Am J Physiol Cell Physiol. 2009;297:C217–25.

16. Geng T, Li P, Okutsu M, Yin X, Kwek J, Zhang M, et al. PGC-1alpha plays a functional role in exercise-induced mitochondrial biogenesis and angiogenesis but not fiber-type transformation in mouse skeletal muscle. Am J Physiol Cell Physiol. 2010;298:C572–9.

17. Leone TC, Lehman JJ, Finck BN, Schaeffer PJ, Wende AR, Boudina S, et al. PGC-1alpha deficiency causes multi-system energy metabolic derangements: muscle dysfunction, abnormal weight control and hepatic steatosis. PLoS Biol. 2005;3:e101.

18. Halling JF, Jessen H, Nohr-Meldgaard J, Buch BT, Christensen NM, Gudiksen A, et al. PGC-1alpha regulates mitochondrial properties beyond biogenesis with aging and exercise training. Am J Physiol Endocrinol Metab. 2019;317:E513–E25.

19. Rowe GC, Patten IS, Zsengeller ZK, El-Khoury R, Okutsu M, Bampoh S, et al. Disconnecting mitochondrial content from respiratory chain capacity in PGC-1-deficient skeletal muscle. Cell Rep. 2013;3:1449–56.

20. Funai K, Summers SA, Rutter J. Reign in the membrane: How common lipids govern mitochondrial function. Curr Opin Cell Biol. 2020;63:162–73.

21. Heden TD, Neufer PD, Funai K. Looking Beyond Structure: Membrane Phospholipids of Skeletal Muscle Mitochondria. Trends Endocrinol Metab. 2016;27:553–62.

22. Mejia EM, Hatch GM. Mitochondrial phospholipids: role in mitochondrial function. J Bioenerg Biomembr. 2016;48:99–112.

23. Heden TD, Johnson JM, Ferrara PJ, Eshima H, Verkerke ARP, Wentzler EJ, et al. Mitochondrial PE potentiates respiratory enzymes to amplify skeletal muscle aerobic capacity. Sci Adv. 2019;5:eaax8352.

24. Senoo N, Miyoshi N, Goto-Inoue N, Minami K, Yoshimura R, Morita A, et al. PGC-1alpha-mediated changes in phospholipid profiles of exercise-trained skeletal muscle. J Lipid Res. 2015;56:2286–96.

25. Lai L, Wang M, Martin OJ, Leone TC, Vega RB, Han X, et al. A role for peroxisome proliferator-activated receptor gamma coactivator 1 (PGC-1) in the regulation of cardiac mitochondrial phospholipid biosynthesis. J Biol Chem. 2014;289:2250–9.

26. Matyash V, Liebisch G, Kurzchalia TV, Shevchenko A, Schwudke D. Lipid extraction by methyl-tert-butyl ether for high-throughput lipidomics. J Lipid Res. 2008;49:1137–46.

27. Ferrara PJ, Lang MJ, Johnson JM, Watanabe S, McLaughlin KL, Maschek JA, et al. Weight loss increases skeletal muscle mitochondrial energy efficiency in obese mice. Life Metab. 2023;2:

28. Lark DS, Torres MJ, Lin CT, Ryan TE, Anderson EJ, Neufer PD. Direct real-time quantification of mitochondrial oxidative phosphorylation efficiency in permeabilized skeletal muscle myofibers. Am J Physiol Cell Physiol. 2016;311:C239–45.

29. Shahtout JL, Eshima H, Ferrara PJ, Maschek JA, Cox JE, Drummond MJ, et al. Inhibition of the skeletal muscle Lands cycle ameliorates weakness induced by physical inactivity. J Cachexia Sarcopenia Muscle. 2024;15:319–30.

30. Prola A, Blondelle J, Vandestienne A, Piquereau J, Denis RGP, Guyot S, et al. Cardiolipin content controls mitochondrial coupling and energetic efficiency in muscle. Sci Adv. 2021;7

31. Summermatter S, Thurnheer R, Santos G, Mosca B, Baum O, Treves S, et al. Remodeling of calcium handling in skeletal muscle through PGC-1alpha: impact on force, fatigability, and fiber type. Am J Physiol Cell Physiol. 2012;302:C88–99.

