## Supplemental Figure S1 and Table S1 for "Skeletal muscle PGC-1α remodels mitochondrial phospholipidome but does not alter energy efficiency for ATP synthesis"

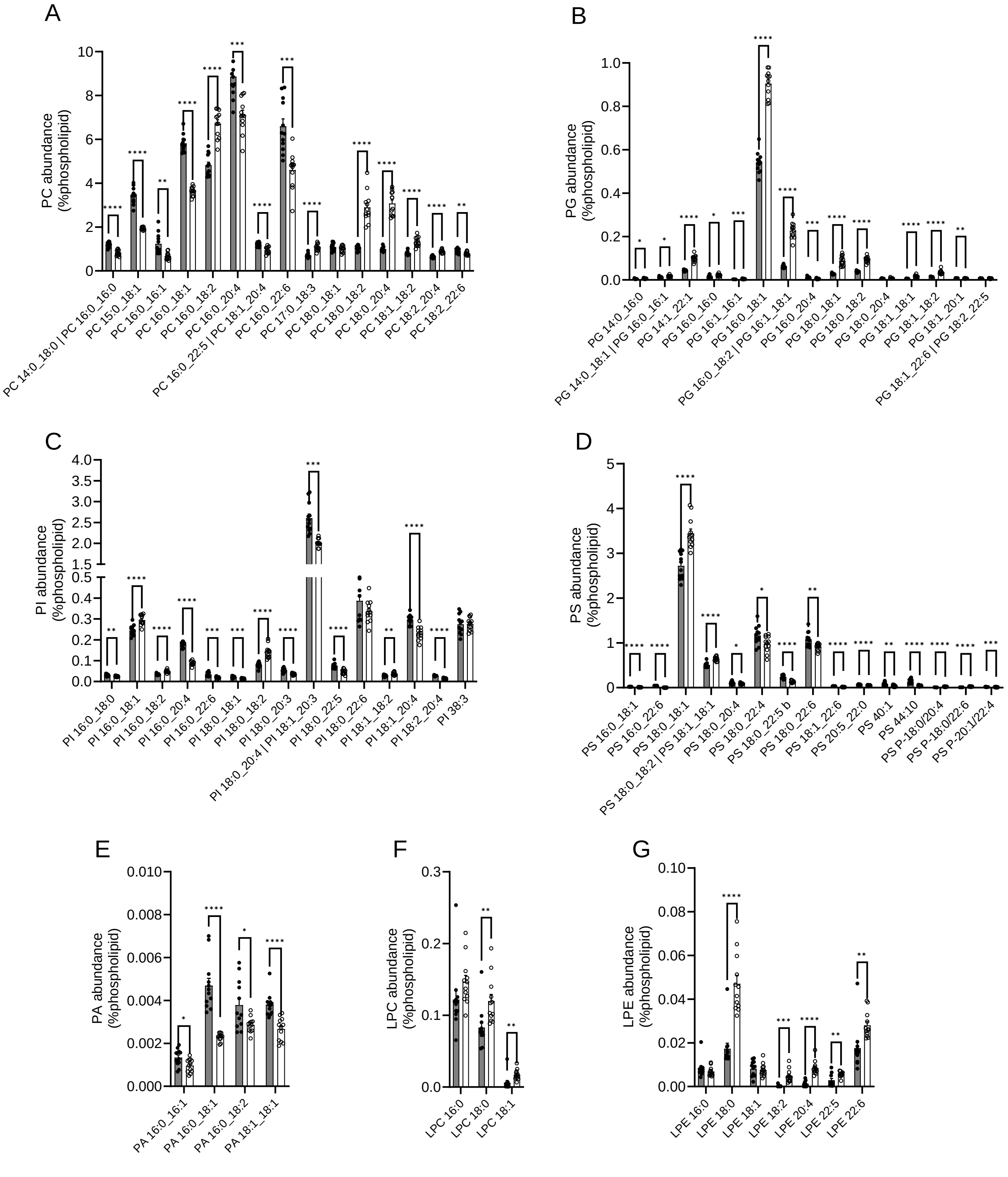
A
PC abundance (%phospholipid)
PC 14:0_18:0 | PC 16:0_16:0
PC 15:0_18:1
PC 16:0_16:1
PC 16:0_18:1
PC 16:0_18:2
PC 16:0_20:4
PC 16:0_22:5 | PC 18:1_20:4
PC 16:0_22:6
PC 17:0_18:3
PC 18:0_18:1
PC 18:0_18:2
PC 18:0_20:4
PC 18:1_18:2
PC 18:2_20:4
PC 18:2_22:6
B
PG abundance (%phospholipid)
PG 14:0_16:0
PG 14:0_18:1 | PG 16:0_16:1
PG 14:1_22:1
PG 16:0_16:0
PG 16:1_16:1
PG 16:0_18:1
PG 16:0_18:2 | PG 16:1_18:1
PG 16:0_20:4
PG 18:0_18:1
PG 18:0_18:2
PG 18:0_20:4
PG 18:1_18:1
PG 18:1_18:2
PG 18:1_20:1
PG 18:1_22:6 | PG 18:2_22:5
C
PI abundance (%phospholipid)
PI 16:0_18:0
PI 16:0_18:1
PI 16:0_18:2
PI 16:0_20:4
PI 16:0_22:6
PI 18:0_18:1
PI 18:0_18:2
PI 18:0_20:3
PI 18:0_20:4 | PI 18:1_20:3
PI 18:0_22:5
PI 18:0_22:6
PI 18:1_18:2
PI 18:1_20:4
PI 18:2_20:4
PI 38:3
D
PS abundance (%phospholipid)
PS 16:0_18:1
PS 16:0_22:6
PS 18:0_18:1
PS 18:0_18:2 | PS 18:1_18:1
PS 18:0_20:4
PS 18:0_22:4
PS 18:0_22:5 b
PS 18:0_22:6
PS 18:1_22:6
PS 20:5_22:0
PS 40:1
PS 44:10
PS P-18:0/20:4
PS P-18:0/22:6
PS P-20:1/22:4
E
PA abundance (%phospholipid)
PA 16:0_16:1
PA 16:0_18:1
PA 16:0_18:2
PA 18:1_18:1
F
LPC abundance (%phospholipid)
LPC 16:0
LPC 18:0
LPC 18:1
G
LPE abundance (%phospholipid)
LPE 16:0
LPE 18:0
LPE 18:1
LPE 18:2
LPE 20:4
LPE 22:5
LPE 22:6

**Fig. S1 Overexpression of PGC-1α remodels mitochondrial phospholipid composition in skeletal muscle** (A–G) Abundance of individual lipid species of phosphatidylcholine (PC) (A), phosphatidylglycerol (PG) (B), phosphatidylinositol (PI) (C), phosphatidylserine (PS) (D), phosphatidic acid (PA) (E), lysophosphatidylcholine (LPC) (F), and lysophosphatidylethanolamine (LPE) (G) in isolated muscle mitochondria (WT n = 12, MCK-PGC-1α n = 12). Data are combined male and female results and are represented as mean ± SEM. * $p<0.05$, ** $p<0.01$, *** $p<0.001$, **** $p<0.0001$ significant difference between groups.

Table S1. Primers for quantitative PCR

| Gene symbol | Forward (5'–3') | Reverse (5'–3') |
|---|---|---|
| Chkb | GCAGAGGTTCAGAAGGGTGA | CCCCAGAAAAAGTGAGATGC |
| Pcyt1 | GGCATGACCAGAGTGAAACA | AGCCCTATGTCAGGGTGACT |
| Pcyt2 | AGGAGAGGTACAAGATGGTACAGG | GCTTCACTTCCTCATAGGTGTCTC |
| Chpt1 | CATCAACCTGGTCACCACAC | CCCAGGGCACATAAAAGGTA |
| Cept1 | TTCTGGAACATTGCGATTTG | AAAAGGTGGTCCTCCAATCA |
| Seleno1 | GCTTTGGATACCAACCCACTCTC | GGTCGAAGTATGTCAGGAGTAGG |
| Pemt | TCCTCAAGGAGTCCAGAGTGAC | ATTGCCACCAGCACCGTCAACA |
| Ptdss1 | ATCACCCTGCTCAGCTTCAC | CAGGATGCCTCTCCAGATGT |
| Ptdss2 | AAACCCCTCAGGATACAGCC | GGAAAATGGCCCGTCTTTAG |
| Prelid3b | GGACTTCGGAGCACGTCTTT | AACTTTCCAGAGGGATCGACA |
| Pisd | GTACAGGGAACGGAAGCTTGA | CCGGAGCCAGTAAGGAAGTT |
| Cds2 | GCTAGATGGAGAGACAGCGT | CGATCATGGCCAAAGTCAGG |
| Prelid1 | CATGACCACCTTCACCTGGAAC | GGATTTCGGTCCAGCCGCTATT |
| Tamm41 | AGACGTGGAAGAGACTCTGCTG | TCTGCCTGATGCTGGAAGGTCT |
| Pgs1 | GAAGTTTCCTTCCGACCTCAAG | AGCAGCATAGTGCGAGAGTTC |
| Ptpmt1 | CTATGAACGAGGAGTACGAGACC | AACTGGACTCCTTTGTGGAGATT |
| Crls1 | TGACCTATGCAGATCTTATTCCA | TGGCAGAGTTCGGTATCTGA |
| Tafazzin | CCCTCCATGTGAAGTGGCCATTCC | TGGTGGTTGGAGACGGTGATAAGG |
| Alcat1 | ATTTTGCTGAGAAGAACGGACTT | TCCACCACAAAGGTAAAGCCA |
| Hadha | TGCATTTGCCGCAGCTTTAC | GTTGGCCCAGATTTCGTTCA |
